# LNP-mediated *BCL11A* Editing Corrects Sickling Phenotypes and Preserves HSPC Fitness Compared to Electroporation

**DOI:** 10.64898/2026.08.26.747413

**Authors:** Yaw Ansong-Ansongton, Corynne S A Adanho, Utku Goreke, Atip Lawanprasert, Mikhail Vysotskiy, York Tang, Alissa LW Kleinhez, Ross Wilson, Angela Rivers, Niren Murthy, David N. Nguyen

**Affiliations:** Department of Bioengineering, University of California, Berkeley, CA; Department of Bioengineering and Therapeutic Sciences, University of California, San Francisco, CA; Innovative Genomics Institute, University of California, Berkeley, CA, USA; Department of Pediatrics, Division of Hematology, University of California, San Francisco, CA; Pioneer Science, Rio de Janeiro, Brazil; Department of Medicine, Division of Hematology/Oncology, University of California, San Francisco, CA; Department of Molecular and Cell Biology, University of California, Berkeley, CA; California Institute for Quantitative Biosciences at University of California, Berkeley, CA, USA; Heritage Medical Research Institute Investigator; Berkeley, CA, USA; Department of Medicine, Division of Infectious Diseases, University of California, San Francisco, CA

**Keywords:** LNP, Electroporation, Cas9, Gene Editing, SCD, RBC Differentiation

## Abstract

Hemoglobinopathies, including sickle cell disease (SCD) and thalassemia syndromes, affect millions of individuals worldwide who have limited access to curative therapies. Autologous hematopoietic stem cell transplant following *ex vivo* CRISPR editing of the *BCL11A* erythroid enhancer reactivates fetal hemoglobin (HbF) and achieves an effective cure, but the resource constraints of clinically approved procedures for editing by electroporation (EP) severely limit widespread implementation. We directly compared the functional outcomes of EP delivery of Cas9 ribonucleoprotein with lipid nanoparticle (LNP) delivery of Cas9 mRNA in primary human HSPCs obtained from healthy HbAA donors and from patients with SCD. While higher editing rates are achieved with EP, LNP-treated HSPCs exhibited greater viability and cell yields that persisted throughout a multi-stage *in vitro* erythroid differentiation protocol. By day 20, the yield of mature red blood cells (CD71^low^CD235a^high^) was lowest in the EP cohorts. Across treatment groups, we observed HbF induction proportional to indel frequency. LNP editing of SCD patientderived HSPCs as low as 25% modified alleles still caused HbF production and reduced the propensity for sickling of *in vitro* differentiated RBCs. These findings highlight the critical trade-offs among manufacturing ease, delivery-associated toxicity, and functional performance across two modalities of therapeutic genome editing for hemoglobinopathies.

**Key points:**

- Although RNP electroporation achieves higher CRISPR editing rates, it reduces HSPC recovery and RBC differentiation; conversely, mRNA-LNP delivery achieves modest rates while preserving HSPC fitness, progenitor expansion, and RBC enucleation efficiencies.
- Despite lower editing rates, mRNA-LNP delivery achieves a sufficient therapeutic threshold to reduce hypoxia-induced sickling without altering bulk oxygen affinity.

## INTRODUCTION

Sickle Cell Disease (SCD) is an autosomal recessive β-hemoglobinopathy caused by a single-base pathogenic variant in the *HBB* gene, resulting in hemoglobin (HbS) polymerization upon deoxygenation, chronic hemolysis, and life-threatening pain episodes.^1–3^ Curative autologous hematopoietic stem cell transplantation (HSCT) has recently been established with CRISPR-Cas9 genome editing disrupting the erythroid-specific enhancer of *BCL11A*.^4–8^ This was recently FDA-approved as exagamglogene autotemcel.^9^ BCL11A acts as a major repressor of γ-globin (*HBG*) expression;^10^ its successful targeting mimics a natural hereditary persistence of fetal hemoglobin with sustained HbF expression, which inhibits HbS polymerization.^11^ *and* can also treat beta thalassemia.

Despite clinical success, current exagamglogene autotemcel manufacturing relies upon electroporation (EP) of Cas9 ribonucleoprotein complexes (protein and guide RNA) into a patient’s HSPCs. This EP process causes cellular toxicity and reduced viability, requires specialized infrastructure, and incurs large startup and operational costs.^12–14^ Lipid nanoparticles (LNPs) represent a potentially scalable alternative for delivering CRISPR cargo, with less centralized infrastructure. Here, we compared *ex vivo* LNP-mediated delivery of Cas9 mRNA with EP delivery of Cas9 protein, both using the same sgRNA targeting the *BCL11A* +58-kb enhancer, in primary human CD34+ HSPCs from non-SCD donors (HbAA) and sickle cell donors (HbSS). Through *in vitro* differentiation, we evaluated editing rates, cellular expansion, functional multipotency, erythroid maturation, HbF induction, oxygen affinity, and protection from hypoxia-induced sickling to define the biological tradeoffs between editing potency and cellular fitness.

## RESULTS

### Genome editing by LNP does not impair HSPC fitness or erythroid potential

The impact of genome editing by electroporation or LNP delivery was assessed across 3 cohorts of experiments (**Table S1**): first in HSPCs from an HbAA healthy donor, then (in the second and third cohorts) in HSPCs obtained from HbSS patients. Following a 48-hour pre-stimulation, HSPCs underwent EP-or LNP-mediated CRISPR editing targeting the *BCL11A* enhancer **(Fig. 1A and B; Table S1)**. Using the Cytiva Life Sciences CD34+ HSC LNP kit on a NanoAssemblr Spark microfluidic mixing platform, we encapsulated Cas9 mRNA and sgRNA with high efficiency **(Table S2)**. At 48 hours post-editing, LNP-treated cells demonstrated greater recovery than EP-treated cells in both HbAA and HbSS HSPC cohorts, with cell recovery numbers for LNP comparable to untreated controls, suggesting minimal toxicity (**Fig. 1C**). We measured editing efficiency using targeted next-generation sequencing to quantify insertions and deletions at the *BCL11A* enhancer region. EP achieved the highest %modified allele, 48hrs post-edit, reaching up to 74%, whereas LNP-mediated delivery achieved up to 41% modified alleles (**Figs. 1D and 1E**).

**Figure 1.**
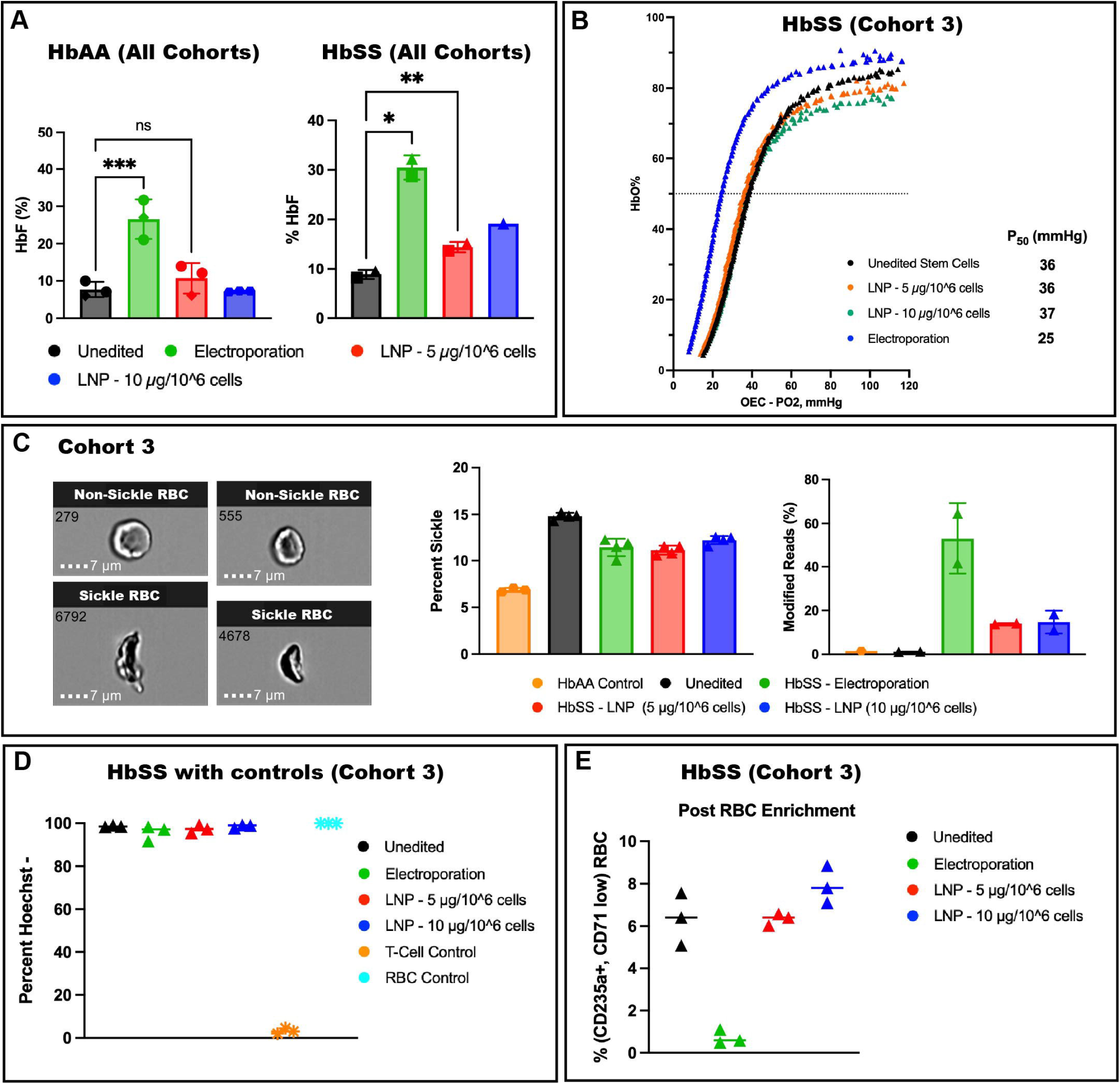
Experimental workflow, HSPC fitness and editing efficiency. (A) Comparison of CRISPR/Cas9 delivery platforms. Detail of the *BCL11A* gene erythroidspecific +58 kb enhancer locus targeted by the specific guide RNA sequence: CTAACAGTTGCTTTTATCAC. The physical electroporation pathway delivers intact Cas9 ribonucleoprotein (RNP) complexes across the cell membrane, whereas the lipid nanoparticle (LNP) pathway encapsulates Cas9 mRNA + sgRNA for endocytic cellular uptake and transient cytoplasmic translation. (B) Schematic of the experimental timeline. Timeline showing human CD34+ HSPC thawing, 48-hour prestimulation, and subsequent CRISPR/Cas9 editing delivery on Day 0 via electroporation or mRNA-LNPs. Downstream testing tracks functional and molecular features across the 20-day differentiation window, including CFU assays, NGS-based editing tracking, flow cytometry, HPLC, oxygen affinity curves, and ImageStream analysis. (C) Post-editing cell recovery data aggregated across all 3 cohorts. Absolute viable cell counts quantified 48 hours post-editing across different delivery conditions for HbAA (left) and HbSS (right) genotypes, including LNP high dose (10 ug/10^6^ cells), LNP low dose (5 ug/10^6^ cells), electroporation, Mock LNP, Mock Electroporation, and unedited stem cells. Data represent mean ± SD from N = 2-3 unique biological donors (HbAA: N = 3; HbSS: N = 2), with each donor shown. Bar colors and legend keys denote experimental treatment conditions. Each unique shape represents an individual biological donor (see Supplemental Table 1 for donor details). (D) Editing efficiency as modified alleles per sample (%) measured by targeted NGS from cohort 1 in HbAA HSPCs (Donor 1) 24 hrs post-editing by electroporation and LNP (5 ug/10^6^ cells), with unedited stem cells, Mock Electroporation, and Mock LNP serving as controls. Bar colors and legend keys denote experimental treatment conditions. Each point represents technical replicates. (E) Editing efficiency as modified alleles per sample (%) measured by targeted NGS from cohort 2 in both HbAA (left, Donor 2) and HbSS (right, Donor 3) HSPCs 24 hrs postediting by electroporation and LNP (5 ug/10^6^ cells), with unedited stem cells serving as controls. Bar colors and legend keys denote experimental treatment conditions. Each point represents technical replicates (n = 2 technical replicates per collection across 2 collections for Donor 3; see Supplemental Table 1 for donor details).

We differentiated HSPCs *ex vivo* into RBCs using a three-phase, 20-day protocol **(Fig. 2A)**. Modified allele frequencies, measured throughout differentiation, peaked by day 9 (EP: 89.6%; LNP: 56.1% in cohort 1), with both LNP-and EP-edited cells exhibiting similar repair outcomes dominated by a +1-bp insertion and a 15-bp deletion (**Fig. S1**). We observed comparable erythroid maturation kinetics and progression of CD71 and CD235a expression across all groups (**Fig. 2A, S2**). Despite initial reductions in cell viability and cumulative yield post-EP, all edited populations exhibited growth kinetics mirroring unedited controls. (**Fig. 2B**). Functional multipotency, measured by a standard methylcellulose colony-forming unit (CFU) assay, was similar between both delivery platforms in HbAA HSPCs. Although edited HbSS samples exhibited reduced total colony formation, the relative distribution of BFU-E, CFU-GM, and CFU-GEMM colonies remained stable regardless of delivery method (**Fig. 2C**).

**Figure 2.**
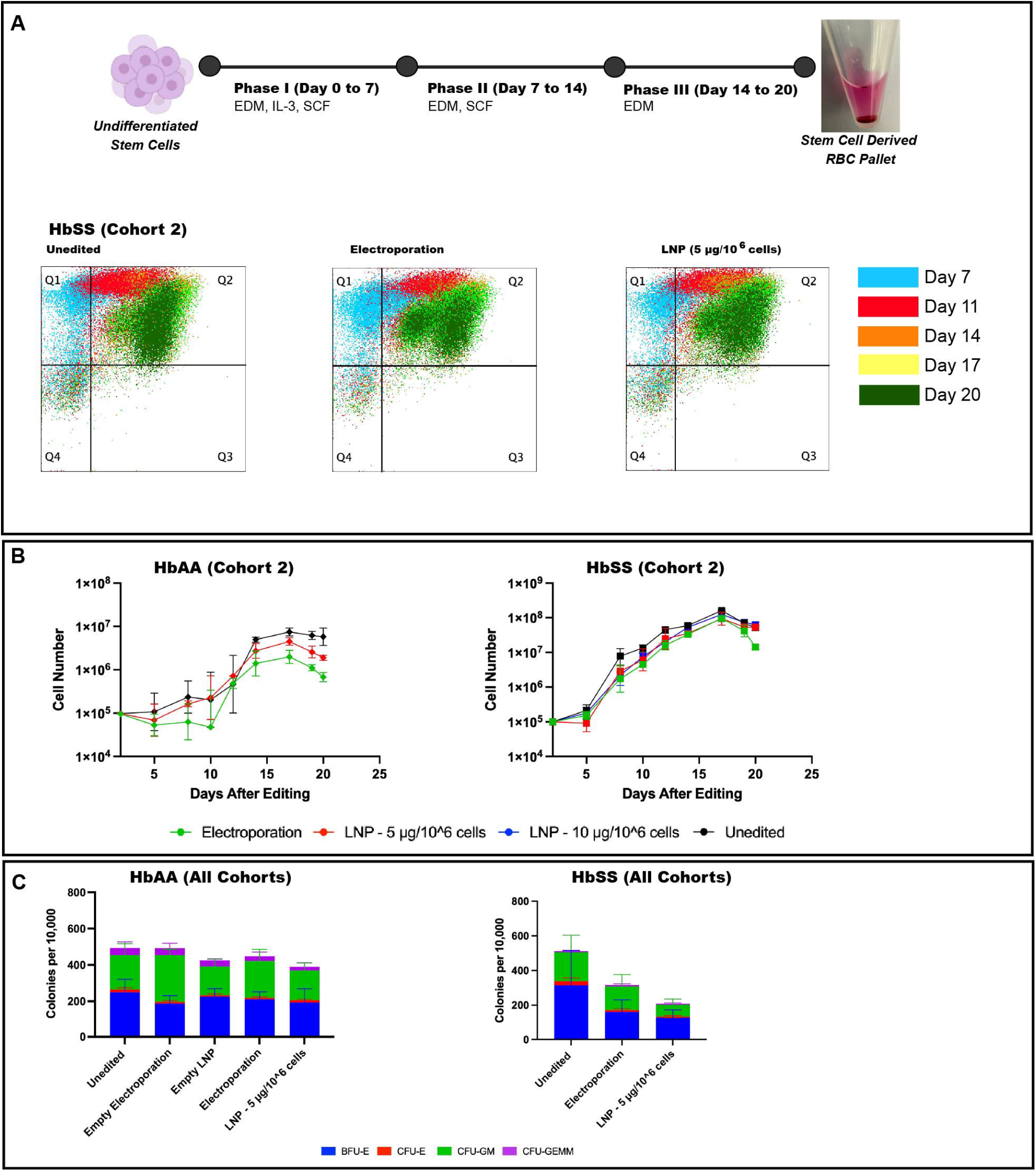
Impact of *in vitro* LNP and electroporation delivery modalities on erythroid differentiation layout, expansion kinetics, and functional multipotency. (A) Schematic flow of the 3-phase liquid culture protocol (Phase I, Day 0 to 7 with EDM, IL-3, and SCF; Phase II, Day 7 to 14 with EDM and SCF; Phase III, Day 14 to 20 with EDM) yielding a stem-cell-derived mature RBC pellet. Representative flow cytometric density plots generated from cohort 2 samples demonstrate progressive progression of LNP-and EP-edited HSPCs from CD71^high^CD235a^low^ erythroid progenitors to mature CD71l^ow^CD235a^high^ erythropoietic cells over 20 days. Untreated HSPCs serve as controls. (B) Representative total cell expansion and viability profiles tracked over 20 days of erythroid culture (showing HbAA Donor 2 and HbSS Donor 1; see Supplemental Table 1). Points represent mean ± SD of independent differentiation replicates for the indicated donors, with error bars between replicates shown. Line colors and legend keys denote experimental treatment conditions. (C) Colony-forming unit (CFU) potential across all cohorts. Absolute distribution profiles of erythroid (BFU-E, CFU-E) and myeloid (CFU-GM, CFU-GEMM) lineages per 10,000 cells seeded in methylcellulose for HbAA (left, N = 3 biological donors) and HbSS (right, N = 2 biological donors) backgrounds. Bars represent mean ± SD across biological donors (see Supplemental Table 1 for donor details).

### *BCL11A* Editing Induces Fetal Hemoglobin and Rescues the Sickle Phenotype

For terminal erythroid products at day 18 of differentiation, EP induced the highest HbF levels (27% HbAA, ∼30% HbSS) measured by HPLC (**Figs. 3A, S3**) and produced the largest increase in oxygen affinity (P50: 25 mmHg) (**Fig. 3B**). However, hypoxiainduced sickling, measured by morphology assessment after timed deoxygenation and fixation (**Figs. 3C, S4**), was improved in both EP-and LNP-edited cells compared with untreated controls. Representative images confirmed preservation of normal erythroid shapes following editing, whereas unedited HbSS cells displayed characteristic sickling under hypoxic stress **(Fig. 3C**). LNP-treated cells had a relatively lower HbF fraction (∼10.7% HbAA, ∼14% HbSS, **Fig. 3A**), yet these levels were sufficient to confer functional improvement in sickling (**Fig. 3C**).

**Figure 3.**
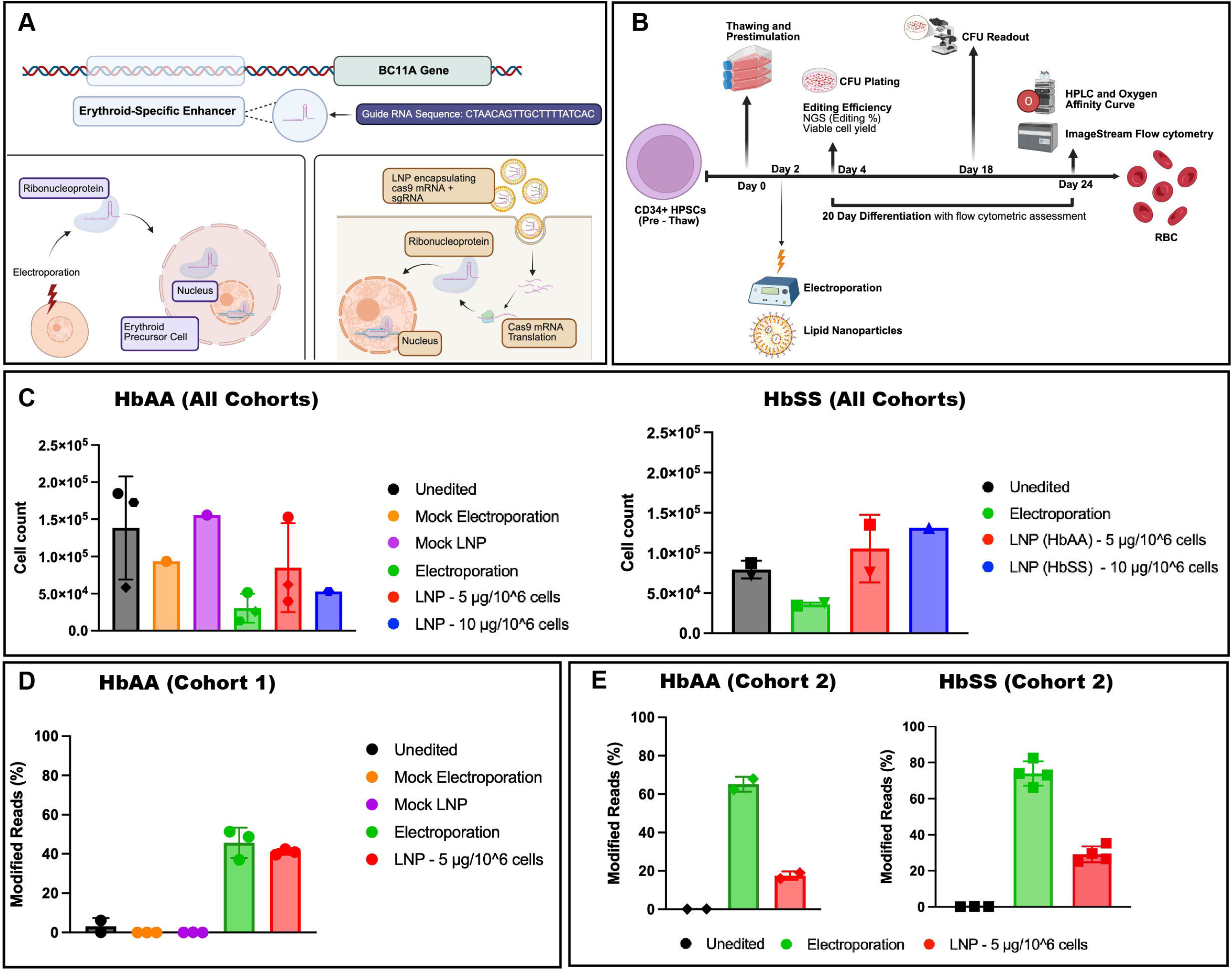
Functional rescue from hypoxia-induced sickling and physiological characterization of terminal erythroid products. (A) Fetal hemoglobin induction data from all cohorts. HPLC-based absolute quantification of HbF percentages (%) achieved in differentiated products from HbAA (left) and HbSS (right). Horizontal significance bars indicate statistical differences compared to the unedited baseline. Data aggregated across 3 cohorts of experiments, with each unique shape indicating a unique biological donor and each point representing technical replicates (HbAA: N=3 across donors; HbSS: N=2 across donors). 3). Bar colors and legend keys denote experimental treatment conditions. Each unique shape represents an individual biological donor (see Supplemental Table 1 for donor details). Statistical indicators: ***P < .001; **P < .01; *P < 0.05; ns, not significant. Statistical analysis was performed using a repeatedmeasures one-way ANOVA with a Geisser-Greenhouse correction, followed by Holm-Šídák’s multiple comparisons test. (B) Oxygen Equilibrium Curves (OEC) from cohort 3 samples. Curve colors and legend keys denote experimental treatment conditions. Continuous functional curves tracing fractional hemoglobin oxygen saturation (HbO%) versus partial pressure of oxygen (OEC - PO2, mmHg). Values for partial pressure at 50% saturation (P50) are summarized for unedited stem cells (36 mmHg), LNP low dose (36 mmHg), LNP high dose (37 mmHg), and electroporation (25 mmHg). (C) Hypoxic sickling rescue from cohort 3 samples. ImageStream-based quantification tracking the percentage of sickled cells under hypoxic stress across groups. Representative multispectral brightfield imagery (left) demonstrates clear biconcave cell morphology in edited or healthy cohorts compared to distorted, sickled morphologies. Scale bars = 7 um. The bar graph (middle), with a baseline detection of ∼6.8% in the HbAA control group, represents the technical limit of detection of the automated morphology gating (e.g., misclassification due to cell orientation/rotation during flow), thereby establishing the functional baseline for zero true sickling in this assay. The bar graph on the right shows the corresponding indel rates for cohort 3 with individual dots representing technical replicates. Bar colors and legend keys denote experimental treatment conditions. (D) Enucleation efficiency data from cohort 3 samples. Percent enucleation was evaluated via Hoechst staining for unedited, electroporated, and LNP conditions relative to non-erythroid T-cell controls and primary mature RBC controls. Each point represents technical replicates. (E) Mature RBC (CD71^low^CD235a^High^) recovery data from cohort 3 samples measured by flow cytometry of HSPCs postfiltration by an Acrodisc™ white blood cell syringe filter. Each point represents technical replicates.

RBCs derived *in vitro* from both LNP-and EP-edited HSPCs progressed through enucleation during differentiation (LNP, 99.2%; EP, 97.2%) (**Fig. 3D, S5**). At day 18, differentiated cells were enriched for RBCs using an Acrodisc™ WBC filter. The input samples had a mature RBC percentage (CD71^low^CD235a^High^) of <2% (**Fig. S6**). Postfiltration, the EP-treated sample had 0.59% mature RBC, but unedited controls and LNP-edited samples had >6% mature RBCs **(Fig. 3E, S6, S7**).

## DISCUSSION

Successful gene editing for SCD depends not only on editing efficiency but also on HSPC fitness and erythroid maturation potential. Our study highlights a trade-off between delivery modalities: electroporation achieved higher indel frequencies but markedly reduced cell recovery, expansion, and mature erythrocyte output. LNPs, however, facilitated HbF induction and functional rescue from sickling, while maintaining cellular expansion and maturation kinetics indistinguishable from unedited controls. This aligns with previous studies comparing EP to other non-viral delivery modalities, including LNPs.^12,14,15^

Interestingly, we observe a disconnect between modified allele frequency and functional outcomes. Despite lower editing rates, LNP-treated cells resisted hypoxia-induced sickling comparably to EP-treated cells. While preferential expansion of edited HSPCs could explain this, both indel rates and erythroid outcomes remained relatively stable throughout differentiation (**Figs. S1, S2**), indicating modification of progenitor cells without specific allele selection or clonal expansion. The editing rates observed in our study compare favorably with established benchmarks under our non-clinical conditions (perhaps limited by using a non-HPLC-purified guide).^16,17^ Although exagamglogene autotemcel reported ∼80–90% edited alleles post-engraftment,^11,18^ the HbF levels we observed fall within clinical amelioration ranges achieved via hydroxyurea, though clinical success depends on total HbF levels and F-cell percentages.^19,20^ Functionally, the HbF induction we achieved with LNPs was sufficient to rescue the sickle phenotype *in vitro*, despite minimal effects on bulk oxygen affinity, supporting the concept that inhibition of HbS polymerization can be achieved independent of oxygen-binding properties.^21^ Given that editing-associated toxicity compromises cellular fitness,^12,22^ achieving a therapeutic threshold of HbF induction may be more important than maximizing editing efficiency.

We acknowledge the lack of F-cell measurements as a limitation, given that the frequency of F-cells clinically predicts therapeutic success perhaps more precisely than bulk HbF measurements.^20,21^ The *ex vivo* differentiation system also lacks the erythroblastic island niche that supports complete *in vivo* RBC maturation.^23,24^ Nuclear extrusion is near-complete (≥ 97%, **Fig. 3D**), but enucleated cells maintained high CD71 surface expression consistent with limitations of a feeder-free *in vitro* erythropoiesis without macrophage-mediated membrane remodeling^30–33^. Still, we observe key characteristics of maturing RBCs with improved function in HbSS-derived cells postgenome editing. Finally, LNP editing at a clinical scale and evaluation in xenograft models will be required to establish impacts on HSPC engraftment and *in vivo* functional improvement.

Overall, these findings support continued development and optimization of mRNA-LNP delivery as an attractive alternative to electroporation for *ex vivo* HSPC editing, particularly for SCD patients unable to mobilize sufficient HSPCs for conventional manufacturing^25^ or those living where HSCT is feasible but clinical electroporation infrastructure is unavailable. By preserving HSPC fitness while achieving sufficient BCL11A disruption to reduce erythroid sickling, LNP-mediated genome editing may provide a more scalable and clinically accessible platform for SCD cure.

## MATERIALS AND METHODS

CD34+ HbAA and HbSS HSPCs were thawed and prestimulated for 24 hours preediting. BCL11A enhancer targeting was performed using either Cas9/sgRNA ribonucleoprotein electroporation or LNP-mediated delivery of Cas9 mRNA and sgRNA.

Cells were subsequently differentiated using a three-phase erythroid differentiation protocol for 20 days, with downstream analysis performed via flow cytometry, CFU, targeted next-generation sequencing, HPLC, and oxygen equilibrium curve analysis. Statistical analyses used one-way ANOVA with Dunnett’s multiple-comparisons testing.

Additional details are in the Supplemental Methods and Figures.

## DATA AVAILABILITY STATEMENT

Data supporting the findings of this study are available from the corresponding author upon reasonable request.

## ACKNOWLEDGEMENTS

The Nguyen Lab was funded by the Innovative Genomics Institute and by a grant from the Heritage Medical Research Institute. D.N.N was supported by NIH grants L40AI140341 and K08AI153767. Y.A.-A. received funding support from the CIRM training program EDUC4-12790 and the Siebel Stem Cell Institute. Research reported in this publication was supported by the National Institute of Diabetes and Digestive and Kidney Diseases of the National Institutes of Health under Award Number TL1DK139565. The content is solely the responsibility of the authors and does not necessarily represent the official views of the National Institutes of Health. Sequencing was performed at the UCSF CAT, supported by UCSF PBBR, RRP IMIA, and NIH 1S10OD028511-01 grants. Special thanks to Eric Soupene for performing oxygen affinity experiments. Special thanks to Agnieszka Czechowicz for mentoring Y.A.-A. on abstract writing through the U2C-TL1 program. This Manuscript was prepared using CURESC_SCD Research Materials obtained from the NHLBI Biologic Specimen and Data Repository Information Coordinating Center and does not necessarily reflect the opinions or views of the CURESC_SCD or the NHLBI.

## AUTHOR CONTRIBUTIONS

Y.A.-A. designed research, performed research, analyzed and interpreted data, and wrote the manuscript. C.A performed research. U.G. performed research and contributed analytical tools. A.L., Y.T, and N.M generated the LNPs for this work. R.W provided vital reagents. A.K. performed research. A.R contributed vital reagents and contributed to the research. D.N. designed research, analyzed and interpreted data, and wrote the manuscript.

## DECLARATION OF INTERESTS

D.N.N. is a member of the scientific advisory board for and owns stock in Navan Technologies.

## Notes

### Summary of Updates

This version of the manuscript has been revised to update the following (400 words maximum; symbols/accents not permitted): 1. Author information has been updated for the corresponding author 2. We removed the manuscript summary box (listing word counts, figure/table/reference counts) as this is not intended to appear in the final manuscript text. No scientific content was altered by this change.

