## Supplemental for "LNP-mediated *BCL11A* Editing Corrects Sickling Phenotypes and Preserves HSPC Fitness Compared to Electroporation"

**SUPPLEMENTAL FIGURES**

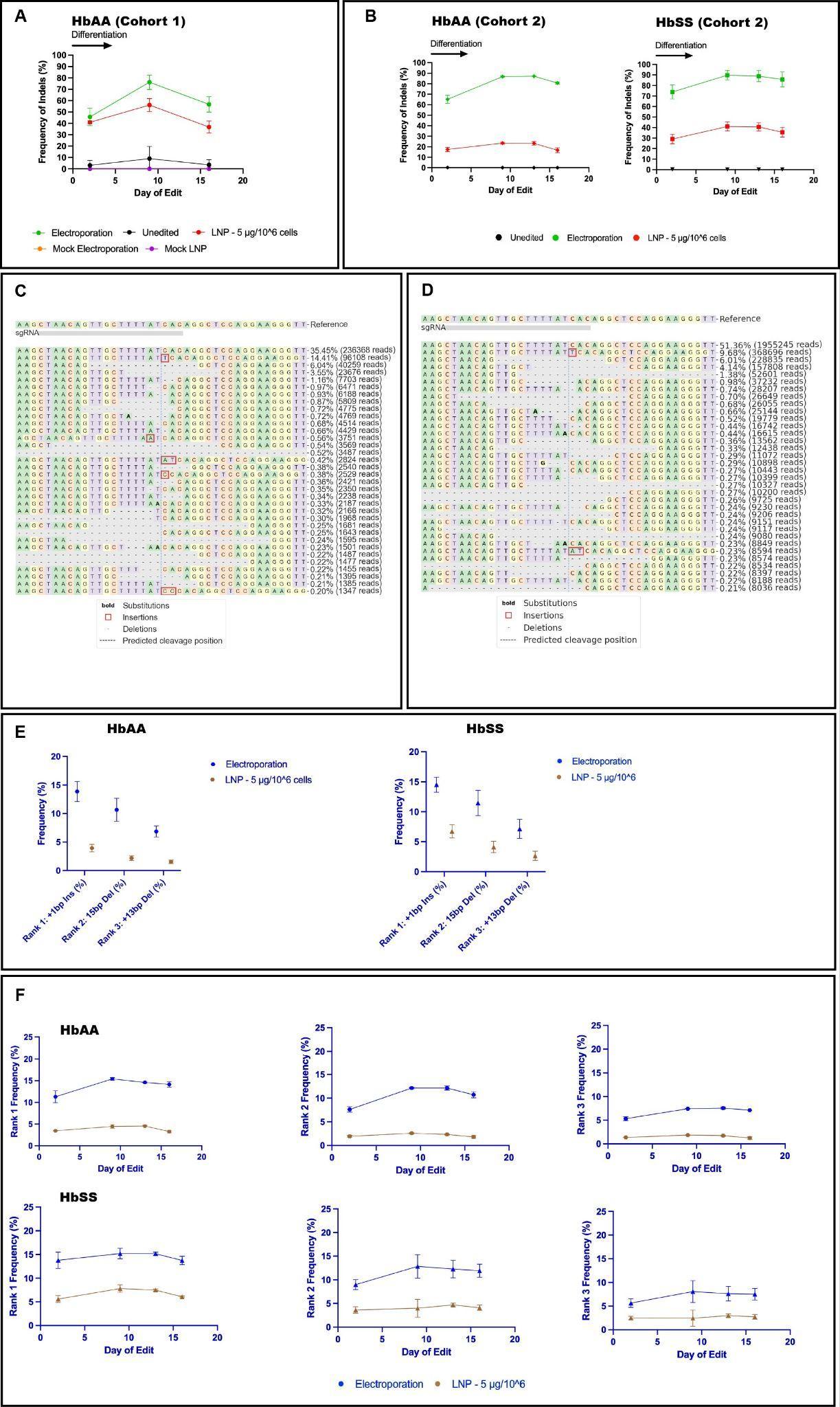

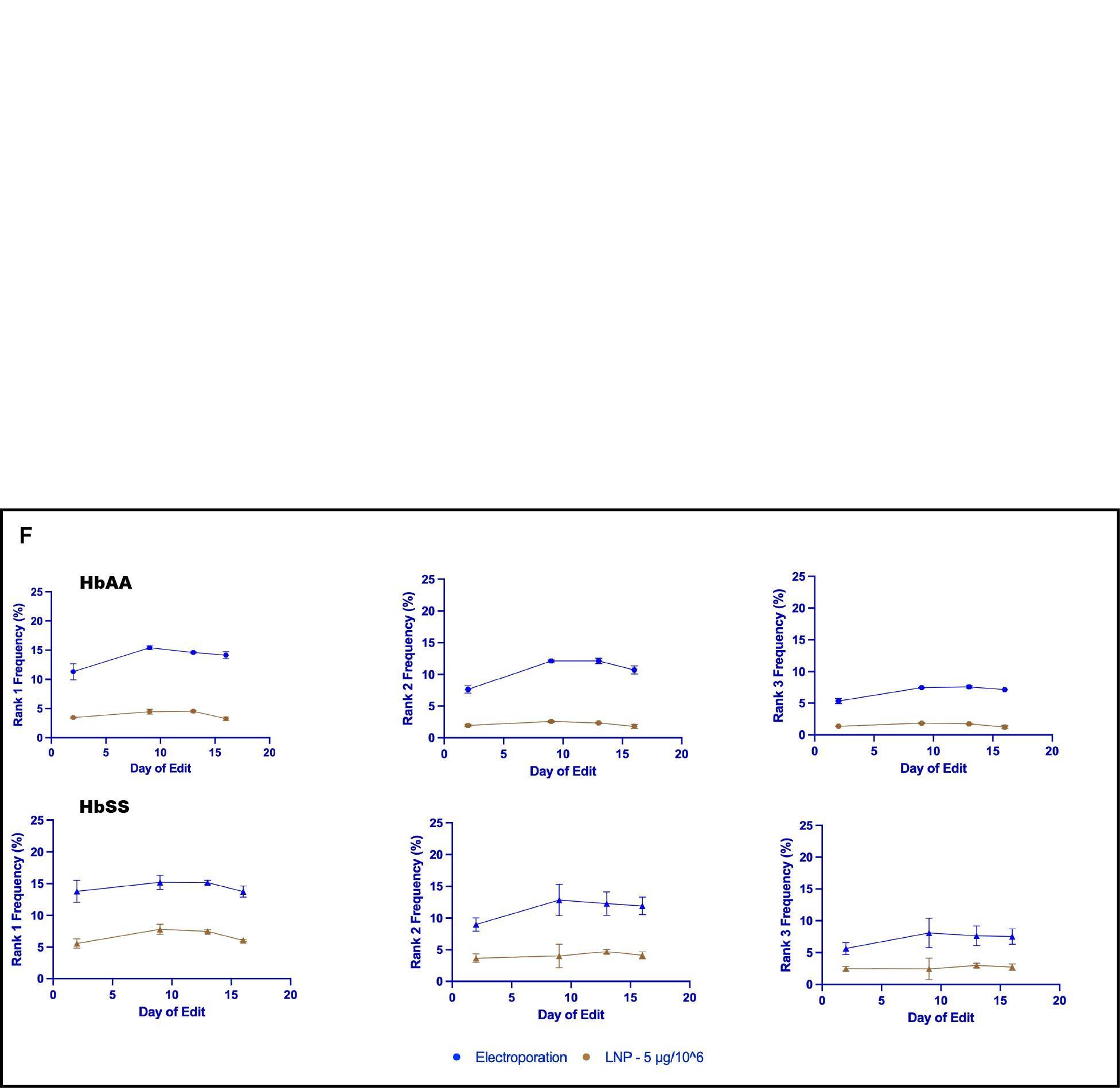

**Supplemental Figure 1. Data showing repair landscape profiles and longitudinal indel stability.** (A) Longitudinal targeted NGS was used to track the frequency of indels (HbAA baseline) across the differentiation timeline for electroporation, unedited stem cells, LNP (5 ug/10^6^ cells), Mock Electroporation, and Mock LNP. Generated from Donor 1. (B) Longitudinal targeted NGS was used to track the frequency of edited alleles in HbAA (left) versus HbSS (right) across the differentiation timeline for electroporation, unedited stem cells, LNP (5 ug/10^6^ cells), Mock Electroporation, and Mock LNP. Generated from Donor 3. (C) Electroporation-mediated (from Donor 1) or (D) LNP-mediated genome editing (from Donor 2) allele frequency profile in primary sickle (HbSS) cells. Genomic DNA was extracted 48 hours post-editing and sequenced with NGS. CRISPResso2 was used to quantify allelic mutation rates. All alleles >0.2% frequency are depicted with alignment to the reference genome at the targeted locus centered around the DSB site, revealing the dominant 1-bp insertion and additional characteristic indels similarly across modalities. (E) Representative frequency of top 3 unique indels across genetic backgrounds and delivery modalities for Rank 1 (+1bp Ins), Rank 2 (15bp Del), and Rank 3 (+13bp Del) side-by-side in HbAA (left) and HbSS (right) patient cells. Data are presented as mean ± SD of samples measured across 3 time points during a 20-day stem cell-to-RBC differentiation protocol. (F) Representative longitudinal tracking of top-ranked alleles through differentiation in HbAA (top) and HbSS (bottom) donors. Data shown are from cohort 1 (top) and cohort 3 (bottom) and are representative of repeated experiments.

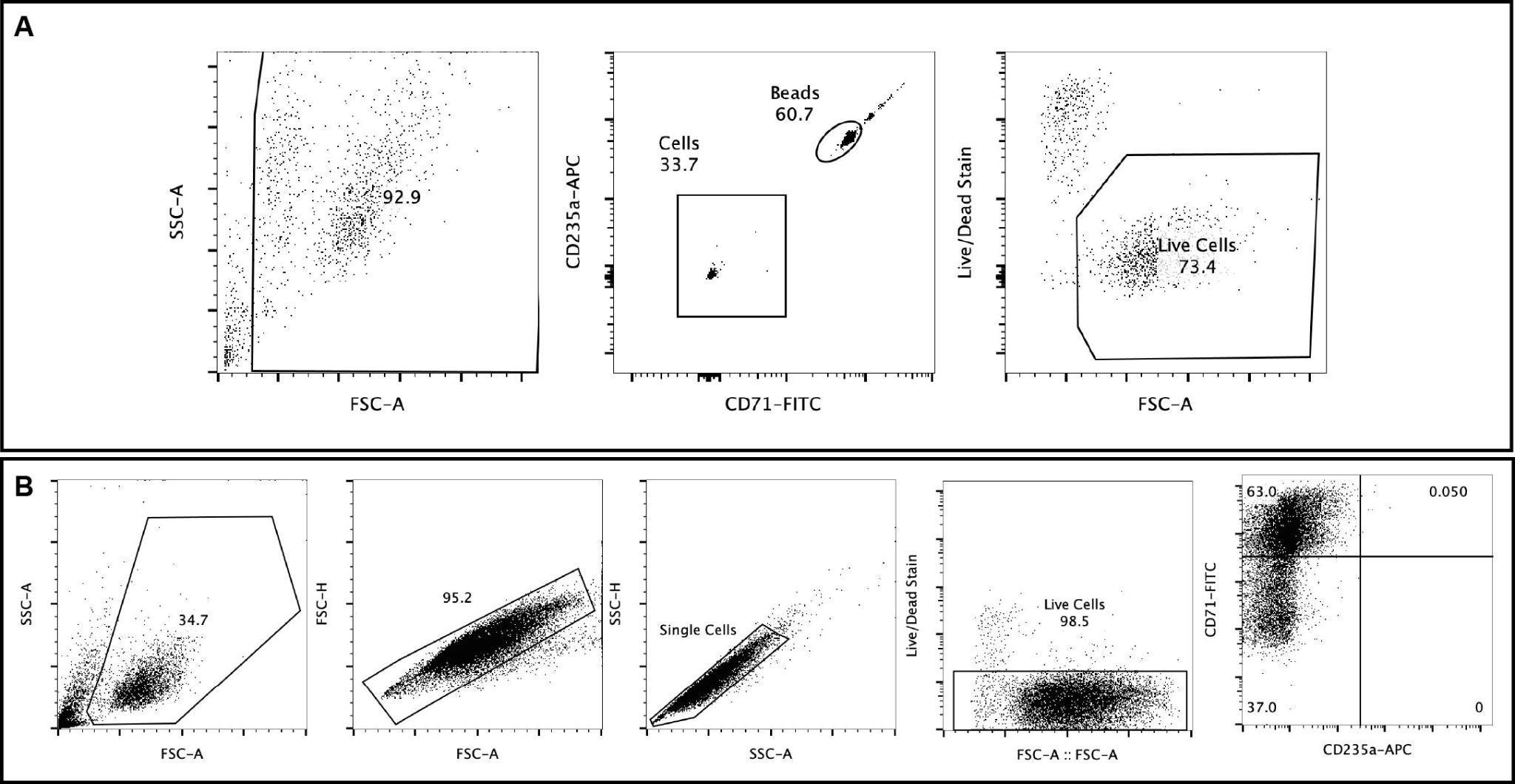

**Supplemental Figure 2. Representative image showing flow cytometry gating strategies for bead-based cell counting and erythroid lineage identification.** (A) Bead-based live cell count strategy. Downstream gating pipeline of live cells utilizing LiveDead/Ghost Dye fluorescent patterns against forward scatter (FSC-A) parameters to ensure absolute cell exclusion and clean isolation of the target live cell populations. (B) Gating strategy to identify mature RBCs based on CD71 and CD235a populations during differentiation. Data shown are from cohort 1 HbSS sample at differentiation day 7 and representative across samples.

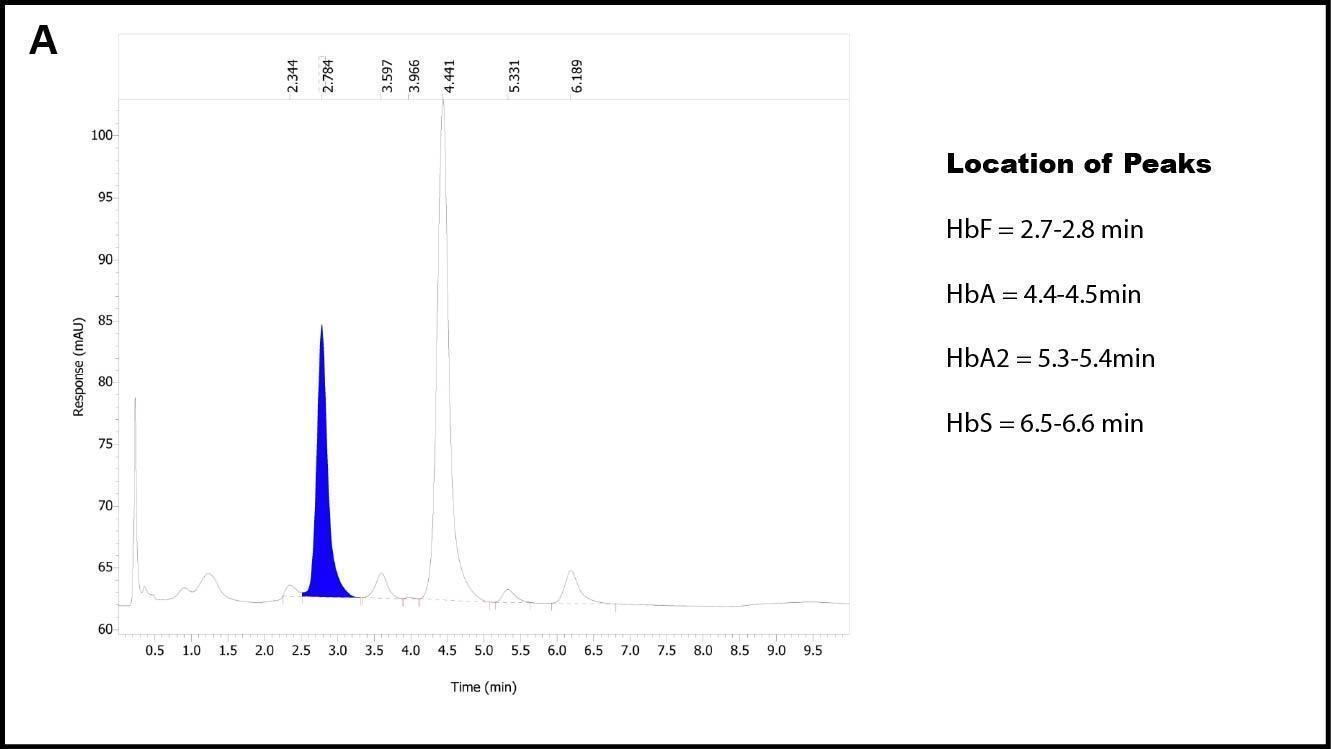

**Supplemental Figure 3. Representative high-performance liquid chromatography trace validation.** (A) HPLC chromatogram trace analysis. Representative milli-Absorbance Units (mAU) were tracked continuously against time (minutes) to validate hemoglobin variant separation. Shaded boundaries indicate separate peak configurations, mapped according to standard reference values: HbF = 2.7-2.8 min (highlighted in blue), HbA = 4.4-4.5 min, HbA2 = 5.3-5.4 min, and HbS = 6.5-6.6 min.

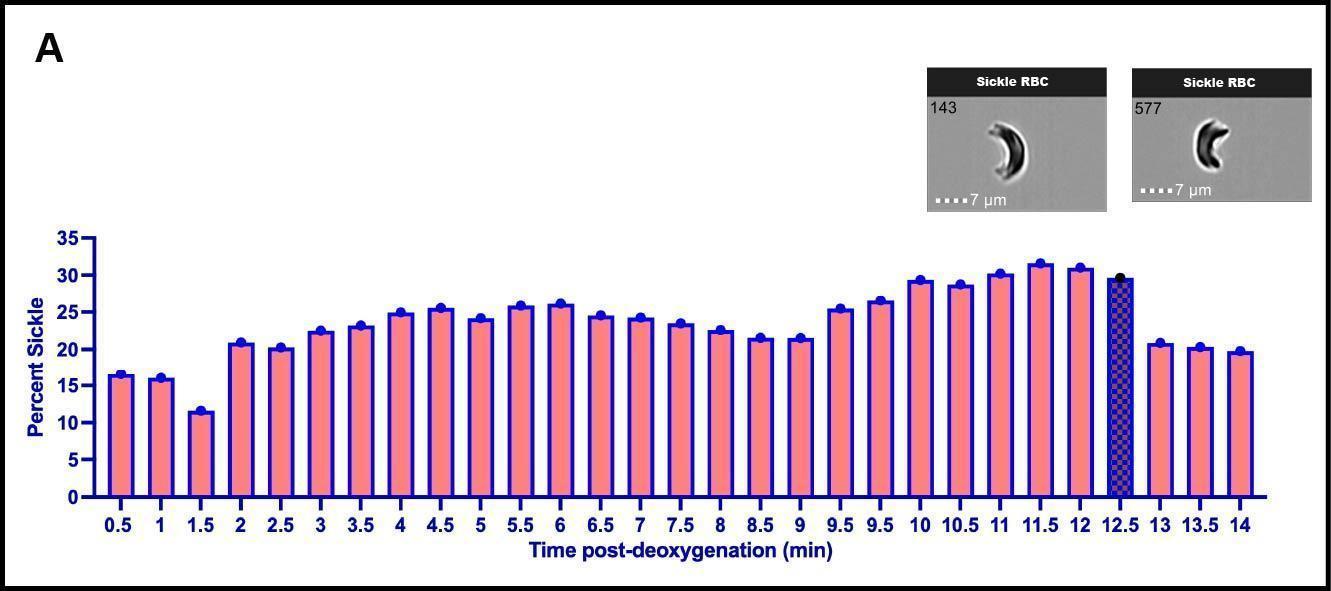

**Supplemental Figure 4. Analytical validation of sickling kinetics and optimal fixation windows.** (A) Time-course of image-based measured sickling profiled for 14min after oxygen chelation with sodium metabisulfite treatment followed by glutaraldehyde fixation. Longitudinal bar graph plotting the measured percentage of sickled red blood cells (%) versus total duration of deoxygenation exposure (minutes). The checkered bar indicates the optimal 12.5-minute stabilization window for sodium metabisulfite treatment prior to glutaraldehyde fixation to maximize the precision of cell imaging in downstream ImageStream workflows. Insets display high-definition brightfield morphological captures of characteristic sickled cells.

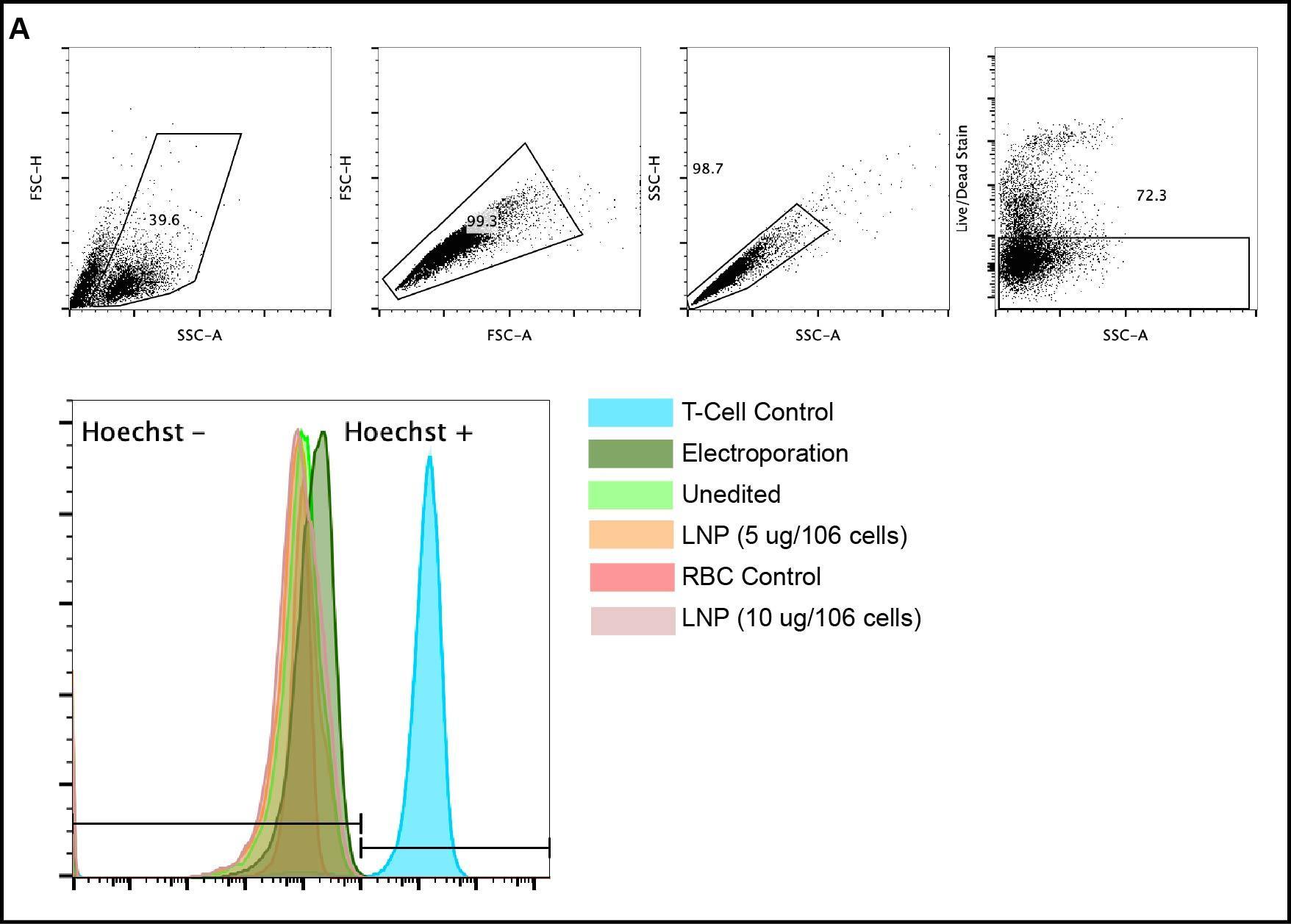

**Supplemental Figure 5. Representative flow cytometry gating strategy for Hoechst stain.** Downstream gating pipeline of live cells utilizing LiveDead/Ghost Dye fluorescent patterns against forward scatter (FSC-A) parameters to ensure absolute cell exclusion and clean isolation of the target live cell populations for onward Hoechst stain.

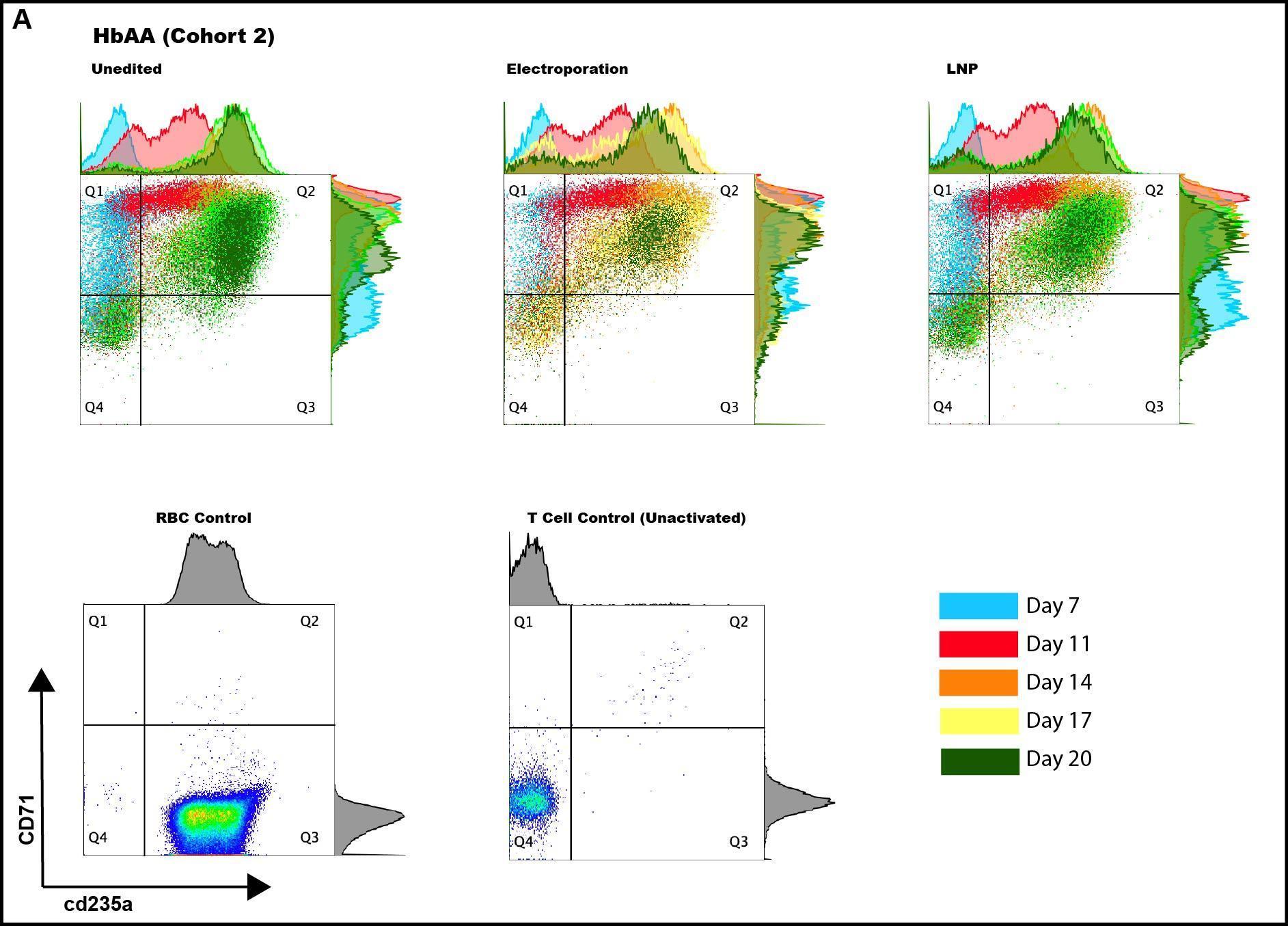

**Supplemental Figure 6. Representative images of comparative flow cytometric profiles of longitudinal erythroid maturation generated from cohort 2 HbAA samples.** (A) High-throughput flow cytometry density plots tracing surface expression profiles of CD71 versus CD235a markers sequentially across culture intervals (Differentiation Day 7, Day 11, Day 14, Day 17, and Day 20). Plots track uniform differentiation-maturation kinetics across Untreated Stem Cells, Electroporated cells, and LNP-edited cell systems for HbAA controls. Baseline references include isolated mature RBC controls and unactivated primary T-cell controls.

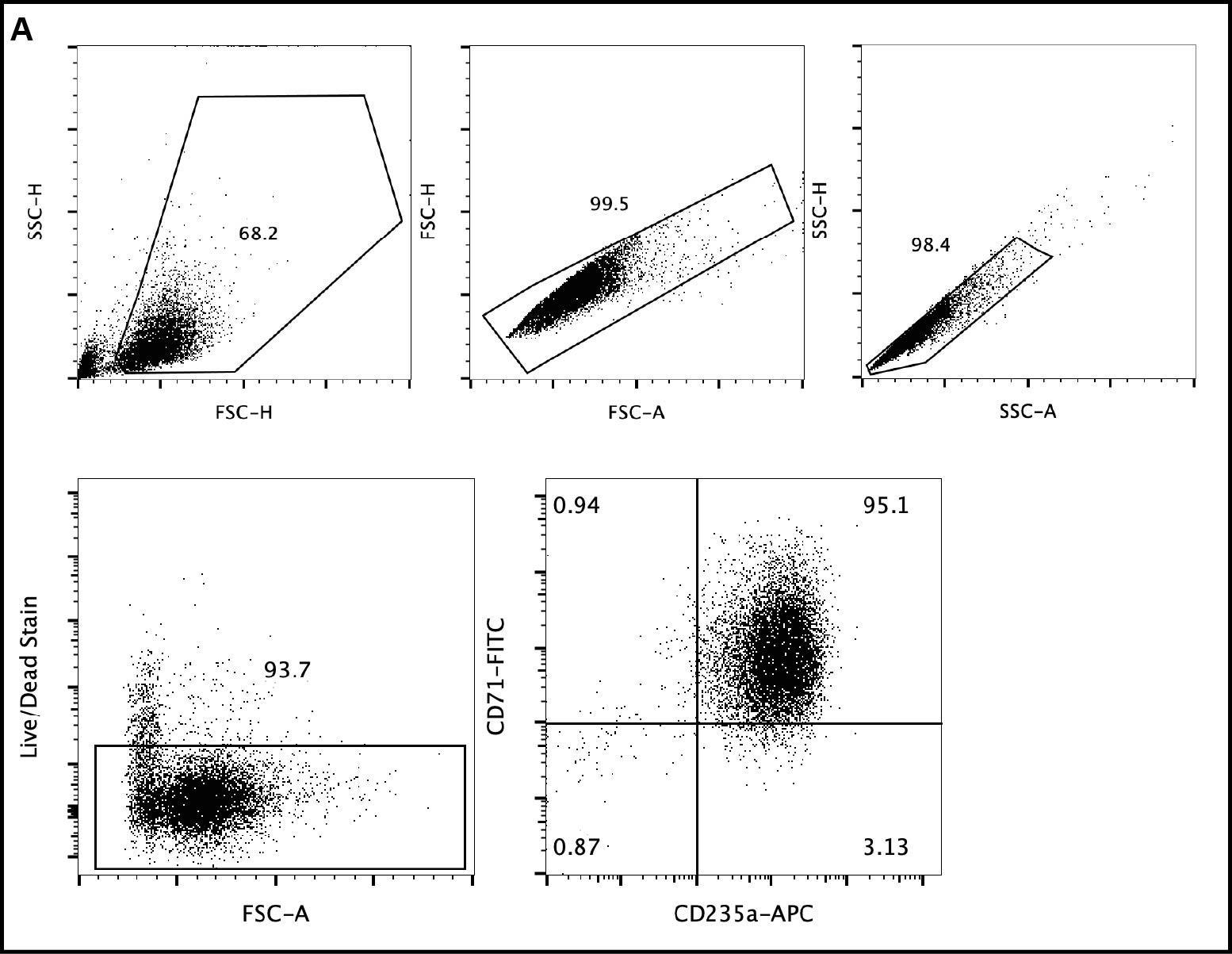

**Supplemental Figure 7. Representative flow cytometry gating strategy for differentiated erythroid cells after filtering through an Acrodisc™ white blood cell syringe filter.** Gating cascades to identify mature RBCs based on CD71 and CD235a populations, alongside Live/Dead populations and forward- and side-scatter profiles.

**SUPPLEMENTAL TABLES**

| **HSPC Donor Number (Symbol)** | **Genotype** | **Figure** | **Source** | **Age (Years) at collection** | **Sex** | **Mobilization method** | **Cohort (Notes)** |
| --- | --- | --- | --- | --- | --- | --- | --- |
| **Donor 1 (⏺)** | HbAA | 1C, 1D, 2C, 3A, S3A | STEMCELL Technologies | 38 | ♀ | G-CSF + Plerixafor | Cohort 1 (N/A) |
| **Donor 2 (⧫)** | HbAA | 1C, 1E, 2B, 2C, 3A, S3B, S7 | STEMCELL Technologies | 26 | ♂︎ | G-CSF + Plerixafor | Cohort 2 (N/A) |
| **Donor 3 (■)** | HbSS | 1C, 2C, 3A, 3B, 3C, 3D, 3E, S3B | BioLINCC | 43 | ♂︎ | Plerixafor Mobilization | Cohort 2  (samples from two collections) |
| **Donor 4 (▲)** | HbSS | 1C, 1E, 2A, 2B, 2C, 3A | BioLINCC | 34 | ♀ | Plerixafor Mobilization | Cohort 3 (N/A) |
| **Donor 5 (⬣)** | HbAA | 1C, 2C, 3A, 3C | AllCells | 30 | ♂︎ | G-CSF | Cohort 3 (N/A) |

**Supplemental Table 1: Key characteristics of HSPCs used across 3 different cohorts**

| **LNP formulation** | **Encapsulation Efficiency (%)** | **RNA Concentration (μg/mL)** |
| --- | --- | --- |
| **Cohort 1** | 81.6 | 53.9 |
| **Cohort 2** | 82.97 | 53.29 |
| **Cohort 3** | 78.97 | 44.65 |

**Supplemental Table 2: Lipid nanoparticle (LNP) formulation quality control attributes.** LNPs encapsulating mRNA and sgRNA were freshly prepared for each discrete editing cohort (Cohorts I-III). Encapsulation efficiency (%) and RNA concentration (μg/mL) are tracked across each formulation.

**SUPPLEMENTAL METHODS**

### **Sample collection**

Peripheral blood samples: This study used de-identified whole-blood samples from volunteer donors with hemoglobin (HbAA), obtained through the Vitalant Denver Blood Donation Center. The original collection protocol was conducted in accordance with the Declaration of Helsinki and approved by the respective Institutional Review Board (IRB) of the collecting site (Vitalant). Vitalant obtained informed consent from all donors prior to blood collection. HbAA samples were collected directly in ACD-B tubes and shipped overnight on ice. The blood samples were centrifuged at 300 G for 5 minutes, and the buffy coat was discarded. The remaining packed red blood cells were then filtered through the Acrodisc™ WBC syringe filters (Pall Corporation) to remove any residual leukocytes. The filtered cells were resuspended at a concentration of 10 million cells/ml in phosphate-buffered saline (PBS) for subsequent experiments.

Stratification by cohorts: Due to the demanding timeline of the longitudinal ex vivo erythroid differentiation protocols, sample processing and treatments were stratified into three independent chronological cohorts (referred to as Cohorts I–III in Supplemental Table 2) with a distinct set of homozygous HbAA and HbSS samples.

CD34 stem cells: Human peripheral blood GCSF-mobilized CD34+ cells (HbAA) were collected from de-identified healthy donors (Allcells). Human peripheral blood GCSF-mobilized CD34+ cells (HbSS) were collected from de-identified sickle cell patients and made available through the NIH NHBLI CURESC_SCD BioLINCC repository. The stem cells were cryopreserved in liquid nitrogen until needed.

**Stem Cell Thawing**
The cryopreserved CD34+ HSPCs were stored in liquid nitrogen (LN_2_) until use. Upon removal from LN_2_, the vial was thawed rapidly at 37 °C. Cells were transferred to a fresh conical tube, then equilibrated by adding 1mL of SFEM II media dropwise while gently mixing, followed by an additional 9 mL of SFEM II to the conical tube. The cell suspension was centrifuged at 100 × g for 10 minutes, and the supernatant was aspirated. The cell pellet was then gently resuspended in 500 μL of media supplemented with DNASE I (final concentration 0.1 mg/mL) (StemCell Technologies Catalog # 07469) and incubated at room temperature for 15 minutes. Following DNase I incubation, the cells were washed in 9 mL of fresh medium and centrifuged again at 100 g for 10 minutes. The supernatant was removed, the cells were resuspended in 1 mL of SFEM II, counted, and finally resuspended in medium with cytokines. The media consisted of 5 mL of SFEM II (Stem Cell Technologies #09605) supplemented with 50 μL of 100x StemSpan CC110 (StemCell Technologies cat #02697) and 50 μL of 100x Penicillin-Streptomycin-Glutamine (ThermoFisher Scientific cat #10378016). The initial seeding density of the cells in media was 450,000 cells per mL.

**Gene Editing**
Electroporation: 24 hours post-thaw, the CD34+ cells were electroporated using the Lonza 4DNucleofector 96-well shuttle system. Ribonucleoprotein (RNP) complexes were assembled at room temperature for 15 minutes by combining the single-guide RNA (sgRNA) (IDT Technologies, Non-HPLC Purified) and Cas9-NLS (40 µM, Berkeley Macrolab). Cas9 was pre-mixed with poly-L-glutamic acid (PGA) (Alamanda Polymers, 25513-46-6) to prevent aggregation before adding the sgRNA.^26^ The final RNP complex contained sgRNA and Cas9 at a 2:1 molar ratio. For electroporation, 200,000 to 300,000 HSPCs were collected per well, centrifuged at 100 × g for 10 minutes, and resuspended in P3 buffer (Lonza Catalog #: V4SP-3096), with 17 μL of cell suspension per reaction. A total of 100pmol (7 μL) of RNP complex was added to the cells, gently mixed, and then transferred to the electroporation plate. Electroporation was carried out with pulse code DS-137. Following electroporation, 80 μL of pre-warmed rescue medium (SFEM II, Stem Cell Technologies #09605, no cytokines) was immediately added. The plate was incubated at 37 °C for 10 minutes, after which the cells were recovered and resuspended in culture media supplemented with cytokines.

Lipid Nanoparticles:

For each cohort, a dedicated microfluidic formulation run was completed on the Cytiva NanoAssemblr Spark platform to generate fresh mRNA-lipid nanoparticles (LNPs), which were then immediately utilized to transfect the corresponding biological donor samples. LNPs carrying Cas9 mRNA and sgRNA were generated using a microfluidic mixing method. The LNP formulation consisted of an aqueous phase containing nuclease-free water, formulation buffer, Cas9 mRNA (Trilink), and sgRNA (IDT Technologies) mixed with the Lipid phase CD34+ HSC Kit (Cytiva cat #29873660) in the NanoAssemblr Spark. All nucleic acid, lipid, and buffer components were prepared at the ratio according to the manufacturer’s protocol. Following formulation, the LNPs were purified using Amicon-100 kDa concentrators into DPBS (pH = 7.4) and quantified for encapsulation efficiency using Ribogreen (ThermoFisher Scientific cat #R11490) before and after addition of 2% Triton-X-100 to disrupt LNPs. For transfection, 200,000 to 300,000 cells were seeded in 6 well Nunc™ Cell-Culture Treated Multidishes (Thermo Scientific, 140675) in 1mL of media consisting of SFEM II medium supplemented with Penicillin-Streptomycin-Glutamine (ThermoFisher Scientific cat #10378016), StemSpan CC110 (StemCell Technologies cat #02697), and 1 μg/mL Gibco™ Human ApoE3 Recombinant Protein (Fisher Scientific cat #35002500UG). The LNPs were administered to the cells at two doses: 5 μg nucleic acid per million cells and 10 μg nucleic acid per million cells.

**Erythroid Differentiation**
Erythroid differentiation was performed in vitro from human CD34+ hematopoietic stem and progenitor cells (HSPCs) using a three-phase liquid culture protocol adapted from the Orkin and Bauer laboratories,^4^ which is based on the foundational method by Giarratana et al.^27^ The differentiation process was conducted in three distinct phases, utilizing Erythroid Differentiation Media (EDM) based on Iscove’s Modified Dulbecco’s Medium (IMDM, Cellgro cat: 15-016-CV) supplemented with 1% L-glutamine (Life Technologies cat: 25030-164) and 2% penicillin-streptomycin (Life Technologies cat: 15140-163). The complete EDM contained Holo-Human Transferrin (330 μg/mL, Sigma cat: T0665), Recombinant Human Insulin (10 μg/mL, Sigma cat: I9278), Heparin (2 IU/mL, Sigma cat: H3149), Inactivated Plasma (5%) (Solvent Detergent Pooled Plasma AB from the Rhode Island Blood Center cat: X0004), and Erythropoietin (3 IU/mL, Amgen cat: NDC 55513-144-10). In Phase 1 (Days 0–7), cells were initially resuspended at a concentration of 5 × 10^4^ cells/mL in EDM I media, which consisted of EDM supplemented with Hydrocortisone (10^-6^ M, Sigma cat: H0135), Stem Cell Factor (SCF, 100 ng/mL, R&D cat: 255-SC), and Interleukin-3 (IL-3, 5 ng/mL, R&D cat: 203-IL). In Phase 2 (Days 7–11), the cells were transitioned to EDM II media and maintained at a density of 2 × 10^5^ cells/mL, consisting of EDM supplemented with only SCF (100 ng/mL). Finally, in Phase 3 (Days 11–18), cells were cultured in EDM III medium at a density of 5 × 10 ^5 cells/mL, with EDM III consisting of the base EDM without additional supplements.

**Next Gen Sequencing (NGS)**
Genomic DNA (gDNA) was harvested at three time points to assess editing efficiency: 72 hours post-thaw (designated as Day 0 of erythroid differentiation), Day 7 of differentiation, and Day 14 of differentiation. For gDNA extraction, harvested cells were centrifuged at 200 g for 10 minutes, after which the medium was aspirated. Cells were washed, transferred to a 0.2mL PCR plate, and the supernatant was aspirated after centrifugation at 400 g for 2 minutes. gDNA was released by adding 50 μL of Quick Extract buffer (Biosearch Technologies #QE09050 to the cells, followed by a 10-minute incubation at room temperature. Lysis was then completed on a thermocycler using the program: 65 °C for 40 minutes, 95 °C for 20 minutes, and a 4 °C hold. The extracted gDNA was used as a template for targeted PCR amplification with PrimeSTAR GXL polymerase (Takara #R050A) using primers specific to the *BCL11A* target locus. The primer sequences used were: Forward (CTCGTTCTGCACATGGAGCTCT) and Reverse (GCAAGAGAAACCATGCACTGGTG). The presence of the amplicon was confirmed on a 1% agarose gel. PCR products were purified for NGS using magnetic beads. The bead solution was added at a 2x ratio of the PCR reaction volume, mixed, and incubated for 10 minutes to facilitate DNA binding. DNA-bound beads were separated on a magnetic rack and washed twice with 180 μL of 70% ethanol. The beads were briefly air-dried. Finally, DNA was eluted with 20 μL of DEPC water. The concentration was confirmed by Picogreen/Tapestation and then indexed for sequencing.

**NGS Analysis**
Targeted deep sequencing data were processed using the CRISPResso2 pipeline (v2.2.9) in a Docker container to quantify *BCL11A* enhancer modification frequencies. See supplementary methods for details.

**Flow Cytometry**
To ensure cellular stabilization and metabolic recovery post-thaw, cells were incubated at 37°C in a 5% CO_2_ humidified incubator for 8 hours prior to immunophenotypic analysis.

Following the recovery period, cells were washed in PBS and resuspended in a staining buffer. Cell viability was assessed using LIVE/DEAD™ Fixable Near IR (780) (ThermoFisher Scientific #L34992). Surface marker staining was performed using a fluorochrome-conjugated antibody cocktail including CD34-PE (BioLegend #378603), CD45-BV421 (BioLegend #304031), and CD38-APC (BioLegend #303509) for 30 minutes at 4°C in the dark. After staining, cells were washed twice and resuspended in PBS + 2% FBS for acquisition.

Cell surface markers were analyzed on Days 7, 11, 14, 17, and 20 of erythroid differentiation. Cells were stained with anti-human CD71-FITC (BioLegend #334104), anti-human CD235a-APC (BioLegend #349113), Hoechst dye (Thermo Fisher Scientific, #33342), and LIVE/DEAD™ Fixable Near IR (780) (ThermoFisher Scientific #L34992) in Dulbecco’s PBS without Ca²⁺/Mg²⁺, 2% FBS, 1 mM EDTA. Cells were incubated at 4°C for 20 minutes in the dark and washed twice before resuspension.

All samples were acquired on a CytoFlex LX (Beckman) flow cytometer and analyzed using FlowJo v10.10.1 software.

**Bead-based Flow cytometric absolute cell counting**

Cell concentrations and viability were determined via flow cytometry using absolute counting beads. Briefly, 10 µL of the unknown cell sample was combined with 10 µL of Precision Count Beads™ (BioLegend, Cat# 424902) and Ghost Dye™ Red 780 (Cytek Biosciences, Cat# 13-0865-T100) in a total staining and reaction volume of 200µL. Following a 20-minute incubation, samples were analyzed on an Attune™ NxT Flow Cytometer with a fixed acquisition volume of 50 µL.

The gating strategy identified the bead population by high fluorescence, while cells were gated by forward and side scatter (FSC-A vs. SSC-A), followed by doublet exclusion and viability assessment (Ghost Dye™-negative). The concentration of the original sample was determined using the ratio of collected cells to beads. To determine the absolute total cell count in the original culture, the calculated concentration was multiplied by the total culture volume (2.0 mL) according to the following formula:

Total cell count = ($\frac{Cell count}{Bead count}x Bead Concentration x Dilution Factor) x Total Media Volume$

In this calculation, a dilution factor of 20 was used to account for the 10 µL sample within the 200 µL reaction volume. The final result represents the total number of viable cells present in the starting experimental volume.

**RBC enrichment**

At the conclusion of the 20-day differentiation protocol (Day 20), mature red blood cells (RBCs) were enriched through the depletion of residual white blood cells (WBCs). The cell suspension was passed through an Acrodisc WBC Syringe Filter (#AP-4952 Cytiva) according to the manufacturer’s instructions.

**Image Stream Flow Cytometry**
To stabilize cell morphology for hypoxic sickling analysis, cells underwent a two-step fixation and stabilization protocol. Briefly, differentiated erythroid cells (enriched for RBC) were treated with 45 mM sodium metabisulfite for 12.5 minutes at room temperature. Following sodium metabisulfite treatment, cells were fixed in 0.05% glutaraldehyde and incubated at 4°C for 48 hours. Samples were then analyzed for sickling morphology using a modified version of the sickling protocol described here.^28^

**CFU Assay**
To assess the clonogenic potential of the cells, a Colony-Forming Unit (CFU) assay was performed. Live cells were counted, and 30,000 cells were resuspended in 1mL of Iscove’s Modified Dulbecco’s Medium (IMDM) supplemented with 2% Fetal Bovine Serum (FBS). This cell/IMDM mixture was then combined with 3 mL of MethoCult^TM^ medium (StemCell Technologies H4434 Classic) by transferring 300 μL of the prepared cell suspension into the methylcellulose medium. The components were mixed thoroughly in a shaker for 5 seconds, then allowed to stand briefly to dissipate any bubbles. The final cell-in-methylcellulose mixture was dispensed into SmartDish^TM^ 6-well plates (#27370, StemCell Technologies) using a 16G blunt-end needle and syringe. To ensure high humidity within the plate, the empty spaces between the wells were filled with sterile H_2_O, and the SmartDish^TM^ plate was then placed into a larger 245mm square dish containing 5mL of sterile water. The assay plates were incubated at 37 °C with 5% CO_2_ for 2 weeks, after which they were inspected under a brightfield microscope for erythroid and myeloid colony formation and quantified.

**Oxygen Affinity**

Oxygen affinity curves were generated as described here.^29^

**HPLC**
HPLC was performed as described here.^16^

**Statistical Analysis**
All statistical analyses were performed using GraphPad Prism (v11.0.1), and data are presented as mean ± standard deviation (SD) derived from at least three independent biological replicates. For comparisons across multiple experimental arms, including post-enrichment RBC recovery (Figure 3A), fetal hemoglobin induction in both HbAA and HbSS backgrounds (Figure 3C), and cellular sickling percentages (Figure 3E), a repeated-measures one-way analysis of variance (ANOVA) with a Geisser-Greenhouse correction was employed to determine overall significance. Upon obtaining significant ANOVA results, a Holm-Šídák multiple comparisons test was performed. Statistical significance was defined as p < 0.05. Significance levels are reported in the figures as follows: ns (not significant, p > 0.05), *p < 0.05, **p < 0.01, ***p < 0.001, and ****p < 0.0001.

#### **Bioinformatic Quantification of Genome Editing and Clonal Dynamics**

##### **CRISPResso2 High-Throughput Sequencing Pipeline**

Quantification of genome editing at the *BCL11A* erythroid-specific enhancer was performed using the **CRISPResso2** software suite. To process multiple experimental replicates across delivery platforms (LNP vs. Electroporation) and longitudinal time points (Days 2, 9, 13, and 16), a batch processing workflow was implemented.

Paired-end FASTQ files were processed using the CRISPRessoBatch command. Amplicons encompassed a 200bp reference sequence centered on the *GATA1* transcription factor binding motif within the erythroid-specific enhancer of the *BCL11A* gene. A ± 5 bp quantification window around the predicted Cas9 cleavage site was used to distinguish CRISPR-induced indels from background sequencing noise.

##### **Downstream Data Processing and Allelic Classification**

Data post-processing was performed in **R (v4.5.2)**. Raw allele frequency tables were imported and categorized based on the following logic derived from the experimental design:

- **Modified vs. Wild-type (WT):** Reads were classified as "WT" if the CRISPResso2 Is_Unedited flag was TRUE. All other reads were categorized as "Modified."
- **Mutation Categorization:** Modified reads were further sub-classified into **Insertions** (n_ins > 0), **Deletions** (n_del > 0), or **Substitutions** (n_mut > 0).
- **Total Editing Efficiency:** Defined as the sum of all "Modified" read percentages within a given sample.

##### **Clonal Ranking and Longitudinal Stability**

To assess the dominance and persistence of specific repair outcomes, alleles were ranked by their relative frequency within the modified population:

- **Top Clone Identification:** For each sample, unique edited sequences were sorted by Reads_Pct. The three most frequent alleles were identified for longitudinal tracking.
- **Comparative Dynamics:** The frequency of these top three ranks was compared between delivery platforms (Electroporation vs. LNP) across all time points to evaluate the consistency of the editing "scar" over time.
- **Indel vs. Substitution Ratio:** The relative proportion of indels (insertions and deletions) versus single-nucleotide substitutions was calculated to determine the structural composition of the edited population.

**SUPPLEMENTAL DISCUSSION**

On day 18 of differentiation, Hoechst staining demonstrated near-complete nuclear extrusion across all conditions (≥ 97%, **Fig. 3D**). However, flow cytometric profiling post-filtration revealed that the vast majority of these enucleated cells maintained high CD71 surface expression (CD71^high^CD235a^high^) rather than a terminally mature CD71l^ow^CD235a^high^ phenotype (**Fig. 3E, S7, S8**).

This discordance between high enucleation efficiency and low frequency of CD71^low^ cells (<2% input, 0.59% EP, 6.4%–6.8% LNP, 6.4% unedited control) reflects a well-documented characteristic of feeder-free *in vitro* erythropoiesis. In the absence of the bone marrow erythroblastic island microenvironment (specifically macrophage-mediated membrane remodeling and exosome vesicle clearance),^30–33^ newly enucleated reticulocytes delay downregulation of CD71. Consequently, Acrodisc™ syringe filtration enriches morphologically enucleated cells (Hoechst negative), but these cells predominantly exist as early enucleated reticulocytes that have not yet fully cleared transferrin receptor (CD71) surface markers.
